# Evolution of SARS-CoV-2 during the first year of the COVID-19 pandemic in Northwestern Argentina

**DOI:** 10.1101/2022.07.08.499374

**Authors:** Romina Zambrana Montaño, Andrés Carlos Alberto Culasso, Franco Fernández, Nathalie Marquez, Humberto Debat, Mariana Salmerón, Ana María Zamora, Gustavo Ruíz de Huidobro, Dardo Costas, Graciela Alabarse, Miguel Alejandro Charre, Ariel David Fridman, Claudia Mamani, Fabiana Vaca, Claudia Maza Diaz, Viviana Raskovsky, Esteban Lavaque, Veronica Lesser, Pamela Cajal, Fernanda Agüero, Cintia Calvente, Carolina Torres, Mariana Viegas

**Affiliations:** Universidad de Buenos Aires, Facultad de Farmacia y Bioquímica, Instituto de Investigaciones en Bacteriología y Virología Molecular (IBaViM), Buenos Aires, Argentina; Consejo Nacional de Investigaciones Científicas y Técnicas (CONICET), Buenos Aires, Argentina; Instituto de Patología Vegetal, Centro de Investigaciones Agropecuarias, Instituto Nacional de Tecnología Agropecuaria (IPAVE-CIAP-INTA), Córdoba, Argentina; Laboratorio de Salud Pública, San Miguel de Tucumán, Tucumán, Argentina; Laboratorio Central de Salud Pública, San Salvador de Jujuy, Jujuy, Argentina; Laboratorio de Virus Respiratorios y Neurovirosis. Hospital Señor del Milagro, Salta capital, Salta, Argentina; Laboratorio de Virología, Hospital de Niños Dr. Ricardo Gutiérrez, CABA, Argentina

**Keywords:** SARS-CoV-2, evolutionary rate, first-wave lineages, COVID-19, Argentina

## Abstract

Studies about the evolution of SARS-CoV-2 lineages in different backgrounds such as naive populations, are still scarce, especially from South America. The aim of this work was to study the introduction and diversification pattern of SARS-CoV-2 during the first year of the COVID-19 pandemic in the Northwestern Argentina (NWA) region and to analyze the evolutionary dynamics of the main lineages found. In this study, we analyzed a total of 260 SARS-CoV-2 whole-genome sequences from Argentina, belonging to the Provinces of Jujuy, Salta and Tucumán, from March 31^st^, 2020, to May 22^nd^, 2021, which covered the full first wave and the early second wave of the COVID-19 pandemic in Argentina. In the first wave, eight lineages were identified: B.1.499 (76.9%), followed by N.5 (10.2%), B.1.1.274 (3.7%), B.1.1.348 (3.7%), B.1 (2.8%), B.1.600 (0.9%), B.1.1.33 (0.9%) and N.3 (0.9%). During the early second wave, the first-wave lineages were displaced by the introduction of variants of concern (VOC) (Alpha, Gamma), or variants of interest (VOI) (Lambda, Zeta, Epsilon) and other lineages with more limited distribution. Phylodynamic analyses of the B.1.499 and N.5, the two most prevalent lineages in NWA, revealed that the substitution rate of lineage N.5 (7.9 × 10^−4^ substitutions per site per year, s/s/y) was a ∼40% faster than that of lineage B.1.499 (5.9 × 10^−4^ s/s/y), although both are in the same order of magnitude than other non-VOC lineages. No mutations associated with a biological characteristic of importance were observed as signatures markers of the phylogenetic groups established in Northwestern Argentina, however, single sequences in non-VOC lineages did present mutations of biological importance or associated with VOCs as sporadic events, showing that many of these mutations could emerge from circulation in the general population. This study contributed to the knowledge about the evolution of SARS-CoV-2 in a pre-vaccination and without post-exposure immunization period.

## 1. Introduction

The COVID-19 pandemic has deeply impacted all populations worldwide, with more than 500 million reported cases so far (Ritchie 2020). South America has reported approximately 9.2% of those cases and presented epidemiological differences in terms of the lineages and variants that were introduced, emerged and circulated in the region, especially during the first and the second wave of the COVID-19 pandemics (Faria *et al*. 2021; Hodcroft 2021; Torres *et al*. 2021).

Molecular surveillance became an invaluable tool to monitor circulating viruses and implement public health policies to prevent the introduction, dispersion, and establishment of viruses with concerning characteristics. However, although there are more than 11 million SARS-CoV-2 genomes submitted to international public databases, only 2.2% belong to South American countries (GISAID, https://www.gisaid.org/), limiting the knowledge about the evolutionary characteristics of the virus in the region.

Diversity and evolution in SARS-CoV-2 are reflected in the description of lineages, variants and clades. On the one hand, the Pango nomenclature system defines “Pango lineages” that are groups of SARS-CoV-2 genome sequences or clusters of infections with shared ancestry (Rambaut *et al*. 2020). Currently, more than 1700 Pango lineages have been defined (https://www.pango.network/). On the other hand, the World Health Organization has defined variants of concern (VOCs) and variants of interest (VOIs) to emphasize priority attention and monitoring of some groups of SARS-CoV-2 sequences (https://www.who.int/en/activities/tracking-SARS-CoV-2-variants/). These variants have been named using Greeks letters, defining VOCs: Alpha, Beta, Gamma, Delta and Omicron, and VOIs: Epsilon, Zeta, Lambda and Mu, among others.

Proyecto PAIS is the inter-institutional federal consortium of SARS-CoV-2 genomics in Argentina, created to monitor SARS-CoV-2 diversity and evolution in the country, including surveillance of SARS-CoV-2 variants of public health interest (Ministerio de Ciencia Tecnología e Innovación 2020). The country reported the first cases of COVID-19 in early March 2020 and experienced a long initial wave from May 2020 up to mid-February 2021, with 2 million cases reported (Ministerio de Salud de la Nación 2020a), in a context of border closures and severe restrictions on internal movement in most of that period (Ministerio de Salud de la Nación 2020b). Subsequently, from mid-February to the end of September 2021, the country suffered an aggressive second wave, reaching more than 5 million reported cases and 115,000 deaths (Ministerio de Salud de la Nación, 2021). The National Vaccination Campaign against COVID-19 in Argentina began at the end of December 2020, for the prioritized population (health professionals and strategic personnel), and since March 2021, it has been gradually extended to the general population (starting with those at high risk for severe disease) (Rearte *et al*. 2022).

Even though millions of sequences have been shared in public databases, studies about the evolution of SARS-CoV-2 lineages in different backgrounds such as naive populations, vaccinees or individuals with post-infection immunity are still scarce, especially from South America. These studies would allow a better understanding of the different scenarios where SARS-CoV-2 can evolve. The aim of this work was to study the introduction and diversification pattern of SARS-CoV-2 during the first year of the COVID-19 pandemic in the Northwestern Argentina region and to analyze the evolutionary dynamics of the main lineages found.

## 2. Material and Methods

### 2.1. Sample collection and sequencing of SARS-CoV-2

In this study, we analyzed a total of 260 SARS-CoV-2 whole-genome sequences from Northwestern Argentina. Among these, 144 sequences of the total dataset correspond to nasopharyngeal swabs samples from patients confirmed for COVID-19 by real-time RT-PCR that showed a cycle threshold (CT) < 30. These samples were randomly selected among the total number of positive cases without an epidemiological link or international travel record, from the Provinces of Jujuy (n= 56), Salta (n= 57) and Tucumán (n= 31).

The period of sampling was from March 31^st^, 2020, to May 22^nd^, 2021 (epidemiological week (EW) 14/2020 to EW 20/2021), which covered the full first wave and the early second wave of the COVID-19 pandemic in Argentina **(Figure 1)**.

**Figure 1.**
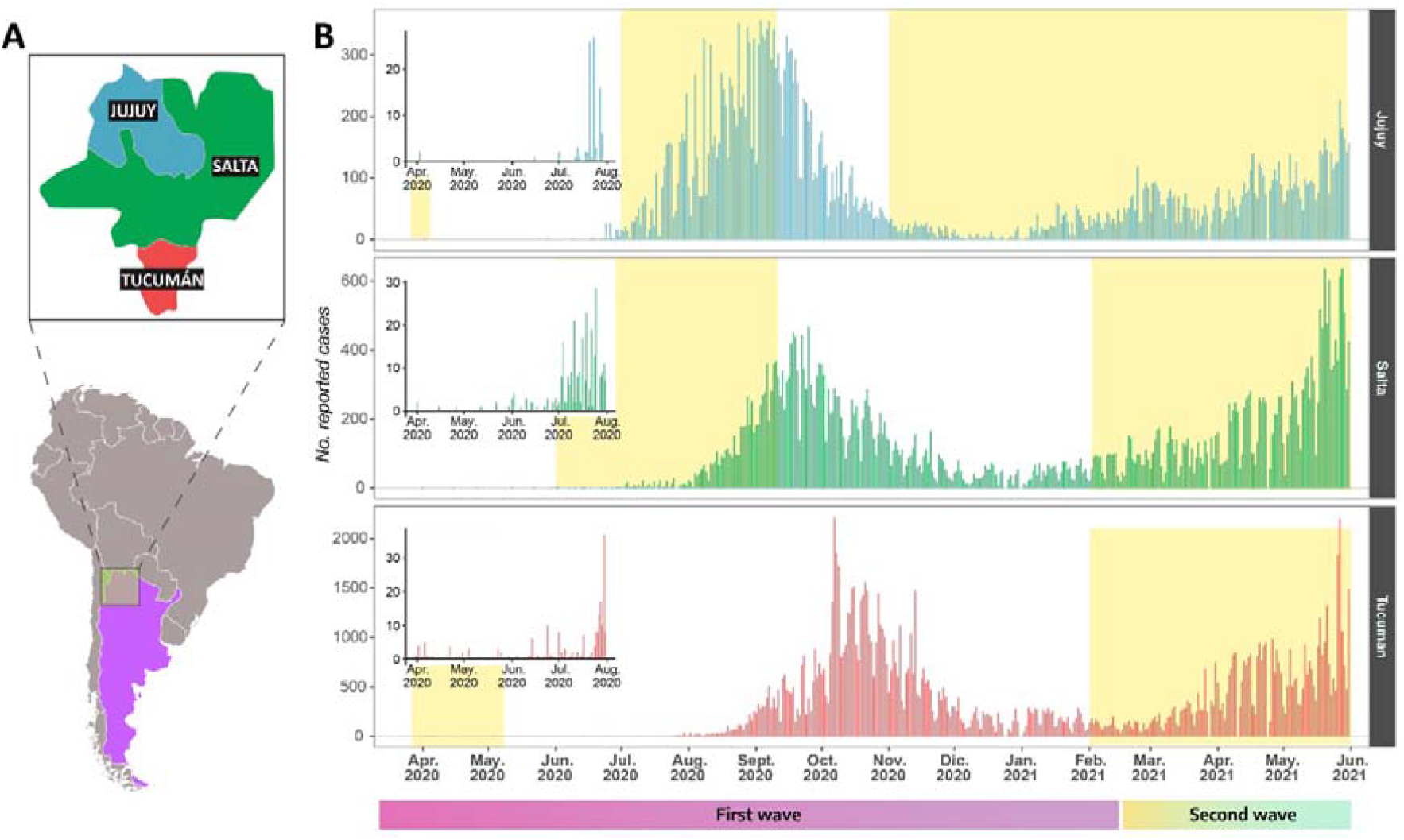
COVID-19 cases diagnosed in Northwest Argentina (NWA) between March 2020 and May 2021. **(A)** Schematic illustration of NWA and its geographic position in the South America map. **(B)** Daily distribution of SARS-CoV-2 cases for studied provinces: Jujuy, Salta and Tucumán. The intervals highlighted in yellow represent the sampling period corresponding to the sequences used in this study (n = 260). The insets show the daily cases for the first five months which had 30 or less daily positive cases.

Nucleic acids were submitted to whole-genome sequencing using the ARTIC protocol with the Oxford Nanopore MinION platform, within the PAIS project.

In addition, sequences from clinical samples were analyzed along with a set of 116 SARS-CoV-2 sequences from samples collected between February and May 2021 in the NWA region, available in the GISAID database (https://www.gisaid.org/) on September 1, 2021, by only selecting complete genomes and excluding those with low coverage: Jujuy (n = 37), Salta (n= 32) y Tucumán (n =47) **(Table S1)**.

### 2.2. Ethical statement

The study was revised and approved by the Medical Ethics and Research Committees of ‘‘Ricardo Gutiérrez’’ Children’s Hospital, Buenos Aires, Argentina (DI-2020-165-GCABA-HGNRG). Informed consent was not obtained, as patient information was anonymized and de-identified before analysis.

### 2.3. Viral classification and assembly of datasets by lineage

SARS-CoV-2 lineages of the sequences obtained in this work were preliminarily assigned according to the Phylogenetic Assignment of Named Global Outbreak LINeages (PANGOLIN) web application (https://pangolin.cog-uk.io/) (Rambaut *et al*. 2020; O’Toole *et al*. 2021). According to the PANGO System (Rambaut *et al*. 2020), seven dominant lineages were determined according to their frequencies in the NWA region: B.1, B.1.1, B.1.499, C.37 (Lambda variant), B.1.1.33, P.1 (Gamma variant), A.2.5.

To separately analyzed the introduction and diversification pattern within the main lineages found, seven datasets were built with the sequences from Argentina, their most similar sequences (the best ten hits for each sequence) from a BLAST analysis conducted against the GISAID database (http://www.gisaid.org, on June 2, 2021) and reference sequences obtained from the PANGO designation list v1.9 (https://github.com/cov-lineages/pango-designation).

We gratefully acknowledge the authors from the originating laboratories responsible for obtaining the specimens and the submitting laboratories where genetic sequence data were generated and shared via the GISAID Initiative, on which part of this research is based (Table S2).

### 2.4. Phylogenetic inference

Sequences for each dataset were aligned using the online version of the MAFFT 7.487 using default parameters (Katoh, Rozewicki and Yamada 2018) (https://mafft.cbrc.jp/alignment/server/), and manually edited in Bioedit version 1.26 (Hall 1999), trimming the first 54 nucleotides of the 5’ end and the last 67 nts of the 3’ end, with respect to the reference hCoV-19/Wuhan/WH04/2020 (EPI_ISL_406801).

Maximum likelihood (ML) phylogenetic trees were built by using IQ-TREE v2.1 (Nguyen *et al*. 2015; Minh *et al*. 2020), employing the best-fit model of nucleotide substitution as selected by ModelFinder (Kalyaanamoorthy *et al*. 2017). The SH-like approximate likelihood-ratio test (1,000 replicates) (Guindon *et al*. 2010) and ultrafast bootstrap approximation (10,000 replicates) (Hoang *et al*. 2018) were used as methods to evaluate the reliability of the groups and branches obtained in trees.

### 2.5. Phylodynamics

Phylodynamic analyses were performed to estimate the time to the most recent common ancestors (ancestral dates of divergence), substitutions rates and demographic reconstructions of the two main lineages found in the NWA region, B.1.499 and N.5. Two datasets including only sequences from the NWA region were compiled.

The temporal signal for both datasets was confirmed using Root-to-tip regression analysis in TempEst v.1.5.3 (Rambaut *et al*. 2016), showing a positive trend of the root-to-tip genetic distances against sampling dates.

The analyses were carried out using an appropriate substitution model for each dataset according to the Bayesian Information Criterion estimated with ModelFInder in IQ-TREE (Kalyaanamoorthy *et al*. 2017; Minh *et al*. 2020). The uncorrelated lognormal (UCLN) molecular clock model (Drummond *et al*. 2006) and the GMRF Bayesian Skyride coalescent model (Minin, Bloomquist and Suchard 2008) implemented in the BEAST v1.10.4 software package, in the CIPRES Science Gateway server (Miller, Pfeiffer and Schwartz 2010) were used. Analyses were run up to convergence, assessed by effective sample size (ESS) values higher than 200 and 10% of the sampling was discarded as burn-in. Uncertainty in parameter estimates was evaluated in the 95% highest posterior density (HPD95%) interval.

### 2.6. Mutation Analysis

To identify which (if any) of the observed mutations was associated with phenotypic changes (such as an alteration of host-cell receptor binding, or changes in viral fitness or antigenicity), all sequences from NWA introduced in this work and from GISAID were analyzed using the CoVsurver tool available at the GISAID EpiCoV platform and the **Outbreak.info** tool (https://outbreak.info/). Three analyses were performed: 1. Genome sequences obtained from the NWA were compared with the Wuhan-Hu-4 reference sequence (GISAID: EPI_ISL_402124), searching for mutations in the Spike gene in the lineages with greater predominance in this study; 2. Mutations of biological importance (mostly characteristic for VOCs) but present in non-VOCs lineages were recorded; 3. Signature mutations for the main monophyletic groups containing Argentinean sequences in different lineages were analyzed.

## 3. Results

### 3.1. Lineage distribution

A total of **16 Pango lineages** from the 260 whole-genome sequences were found in Northwestern Argentina: a region covered by the Provinces of Jujuy (n=93), Salta (n=89), and Tucumán (n=78).

**In the first wave** (from March 2020 to February 2021), eight lineages were identified from the 108 SARS-CoV-2 genomes analyzed: the most predominant one was B.1.499 (76.9%) in the three NWA provinces, followed by N.5 (10.2%), B.1.1.274 (3.7%), B.1.1.348 (3.7%), B.1 (2.8%), B.1.600 (0.9%), B.1.1.33 (0.9%) and N.3 (0.9%).

Instead, **in the early second wave** (March - May 2021), an increase in the lineage diversity was observed but also a different predominant lineage in each province. We found **12 lineages** for the 152 sequences: N.5 (27.6%), P.1 (25.7%), C.37 (11.8%), B.1.1.348 (9.9%), A.2.5 (5.9%), B.1.427 (5.9%), B.1.1.7 (4.6%), B.1.499 (2.6%), P.2 (2.6%), B.1.1.274 (2.0%), B.1.1.519 (0.7%) and C.26 (0.7%).

Most of the sequences from the Provinces of Jujuy and Salta were assigned to the lineage N.5 (n=22/53 and n=15/53, respectively). Lineage P.1 (VOC Gamma, n= 28/46) showed a marked predominance in Tucumán. In addition, the second most frequent lineage was also different for each province: B.1.1.348 (n= 14/53) in Jujuy; C.37 (VOI Lambda, n= 10/53) in Salta and B.1.427 (VOI Epsilon, n= 9/46) in Tucumán **(Figure 2.A)**.

**Figure 2.**
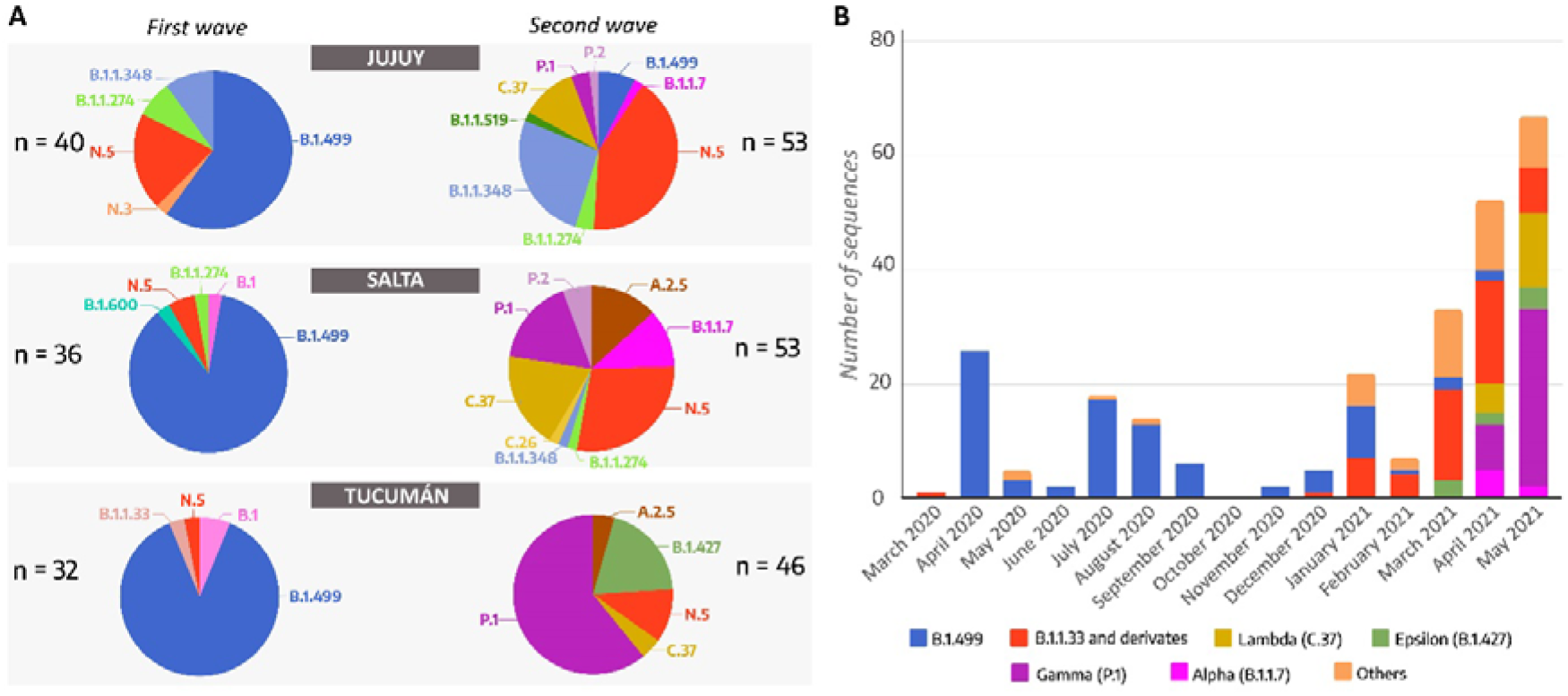
SARS-CoV-2 lineage distribution in Northwest Argentina during the first and second waves. **(A)** Pie charts showing and comparing the proportion of lineages that were detected in the Provinces of Jujuy, Salta, and Tucumán during the first wave and the early second wave in Argentina. **(B)** Temporal representation of SARS-CoV-2 sequences and PANGO lineages in Northwestern Argentina provinces between March 2020 and May 2021.

### 3.2. Spread of the Argentinean lineages B.1.499 and N.5 during the first wave and the early second wave

Argentinean sequences were analyzed along with the most similar sequences in the database to improve the description of monophyletic groups in the NWA, which allow inferring common transmission chains and the establishment of circulating viruses.

**Lineage B.1.499** was initially detected in April 2020 and was the most frequently detected lineage of the first wave in the NWA region. This lineage was introduced at least eight times in this region, originating six clades and two sporadic cases. Three introductions corresponded to monophyletic groups with sequences from the Provinces of Jujuy and Salta, which suggested common transmission chains between these provinces during the first wave **(Figure 3, Clades A - C)**. Furthermore, another group **(Figure 3, Clade E)**, composed of 12 sequences, corresponded to an introduction in the province of Salta that spread to different cities within the province between June and September 2020.

**Figure 3.**
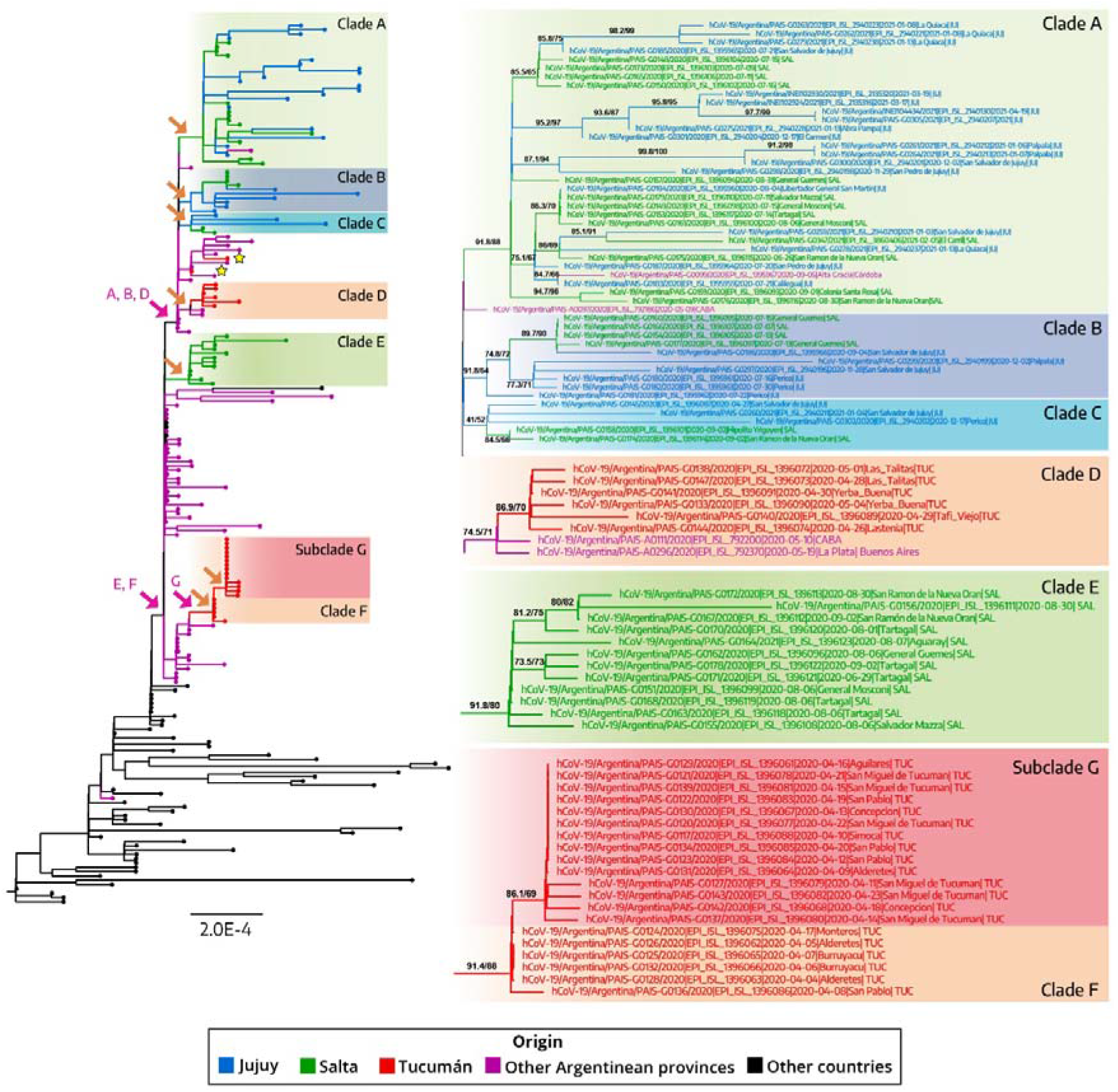
Phylogenetic tree of SARS-CoV-2 whole-genome sequences of lineage B.1.499. The tree was rooted between lineages A and B. Only groups with Northwestern Argentinean sequences are shown and indicated by orange arrows. The stars on the main tree indicate sporadic cases. The SH-like/UFB values for the relevant groups are indicated. Branches and tips are coloured by province. The scale indicates the number of substitutions per site. The pink arrows indicate the context groups used to analyse the descriptive mutations of some clusters that are detailed in **Table 2**.

In contrast, sequences from the Province of Tucumán did not show a phylogenetic association with sequences from the other provinces of the NWA region (Jujuy and Salta), instead, they were more related to sequences from the capital city (Autonomous City of Buenos Aires, CABA) and other provinces located in the Central region of the country, such as Buenos Aires and Córdoba. Four introductions to Tucumán were observed: two corresponded to sporadic cases and the other two involved local groups with viral spread and diversification to different cities of Tucumán during April and May 2020 **(Figure 3, Clades D and F)**.

**Lineage N.5** is an Argentinean lineage (derived from the Brazilian lineage B.1.1.33) that replaced lineage B.1.499 at the end of the first wave (early 2021) **(Figure 2.A)**. During the first wave, at least six introductions in the NWA region were observed, three of them occurred in the Province of Jujuy (January 2021), and only one introduction was observed for Salta and Tucumán (February 2021) **(Figure 4, Clades H – J)**.

**Figure 4.**
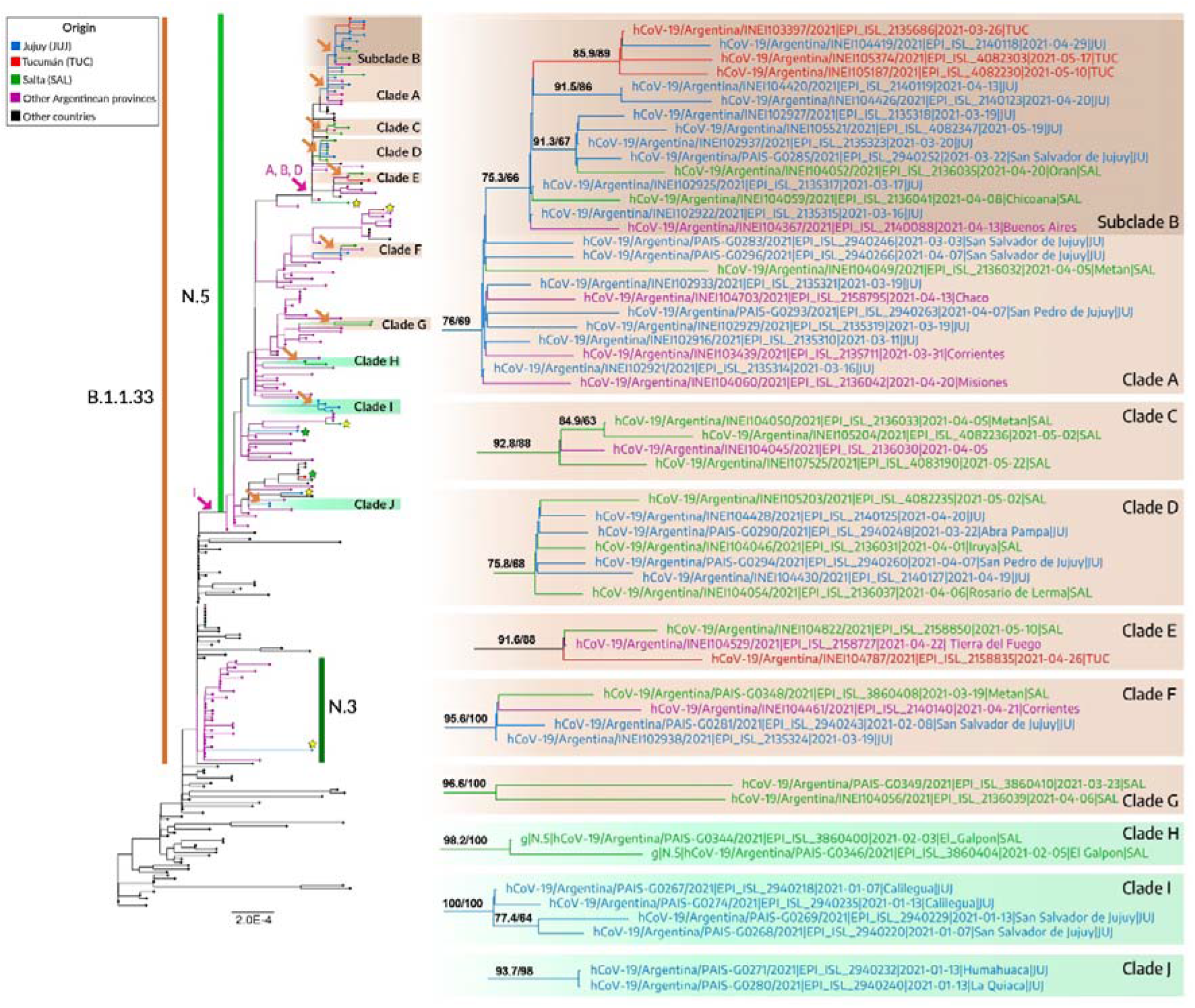
Phylogenetic tree of SARS-CoV-2 whole-genome sequences of lineage N.5. Tree was rooted between lineages A and B. Only groups with Northwestern Argentinean sequences are shown and indicated by orange arrows. The green stars on the main tree indicate sporadic cases detected in the first wave and the yellow stars correspond to the second wave. The SH-like/UFB values for the relevant groups are indicated. Branches and tips are coloured by province. The scale indicates the number of substitutions per site. The pink arrows indicate the context groups used to analyse the descriptive mutations of some clusters that are detailed in **Table 2**. The green panels (H - I) correspond to the first wave and the brown panels (A - G) to the early second wave.

This lineage predominated during the beginning of the second wave in NWA, in March 2021 **(Figure 2)**. However, several introductions of lineages P.1 (VOC Gamma) and C.37 (VOI Lambda) were observed since April 2021, increasing their frequencies in the following month.

Phylogenetic analysis of lineage N.5 showed an increase in the number of introductions in the NWA region as well as in the internal clustering, especially in the Provinces of Jujuy and Salta, which suggested transmission chains between these two provinces. Most of the sequences (n= 16) from Jujuy were grouped in a large clade (Figure 4, clade A) with sequences from Salta (n= 3) and Tucumán (n= 3). In this clade, an inner group was observed (Figure 4, subclade B) with sequences from the three NWA provinces, showing the internal diversification and dispersion of the lineage in the region between March and May 2021.

Few sporadic cases unrelated to other sequences in the NWA region were found in the three provinces: two introductions in the Province of Salta, one in Jujuy and another in the province of Tucumán **(Figure 4)**.

### 3.3. VOC and VOI associated lineages during the early second wave

#### VOC Gamma (lineage P.1)

At the beginning of the second wave, 15 introductions were observed: seven in Salta, seven in Tucumán and one in Jujuy. In general, the introductions of the Gamma variant corresponded to sporadic cases except for a large clade **(Figure 5, clade A)** containing 18 genomes from the Province of Tucuman obtained between April and May 2021. This large clade also contained two sequences from Salta and Jujuy, and a cluster from Chile, evidencing the establishment of the lineage in the NWA region, mainly in the province of Tucumán and its association with circulation in the neighboring country of Chile.

**Figure 5.**
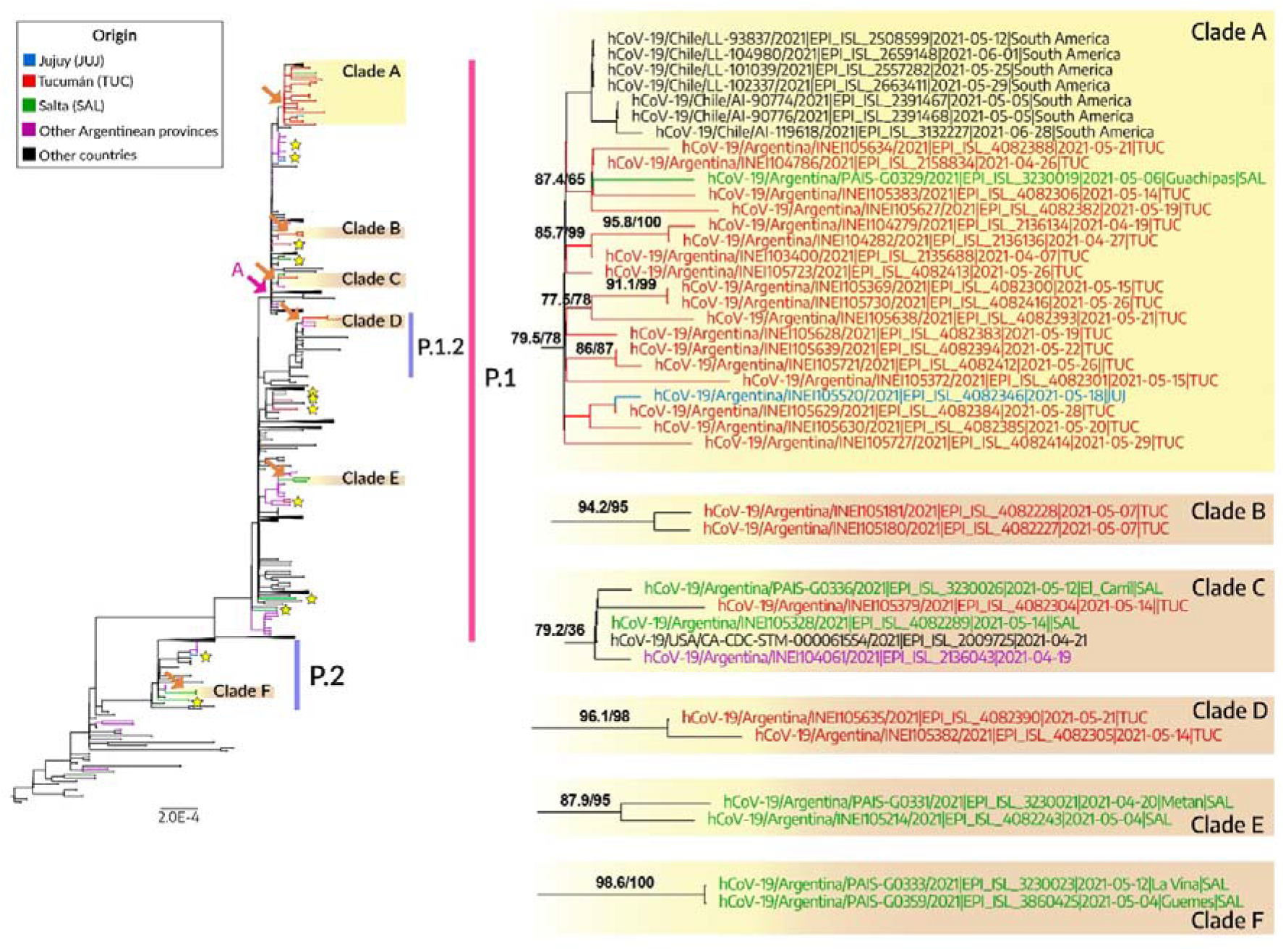
Phylogenetic tree of SARS-CoV-2 whole-genome sequences of Gamma (lineage P.1) and Zeta (lineage P.2). Tree was rooted between lineages A and B. Only groups with Northwestern Argentinean sequences are shown and indicated by orange arrows. The stars on the main tree indicate sporadic cases. The SH-like/UFB values for the relevant groups are indicated. Branches and tips are coloured by province. The scale indicates the number of substitutions per site. The pink arrows indicate the context groups used to analyse the descriptive mutations of some clusters that are detailed in **Table 2**.

Additionally, two sequences collected in May 2021 from Tucumán were associated with the P.1.2 sublineage **(Figure 5, clade D)**.

#### VOC Alpha (lineage B.1.1.7)

Four introductions were observed for this variant, but only one introduction corresponded to a monophyletic group of sequences from Salta (Figure 6, clade A). The remaining samples corresponded to sporadic cases: one from Salta that was associated with sequences from Europe, and two other samples from Salta and Jujuy grouped within a cluster that mostly contained sequences from Argentina. This result showed a brief local transmission period of the Alpha variant during the early second wave, but without establishment in the NWA region.

**Figure 6.**
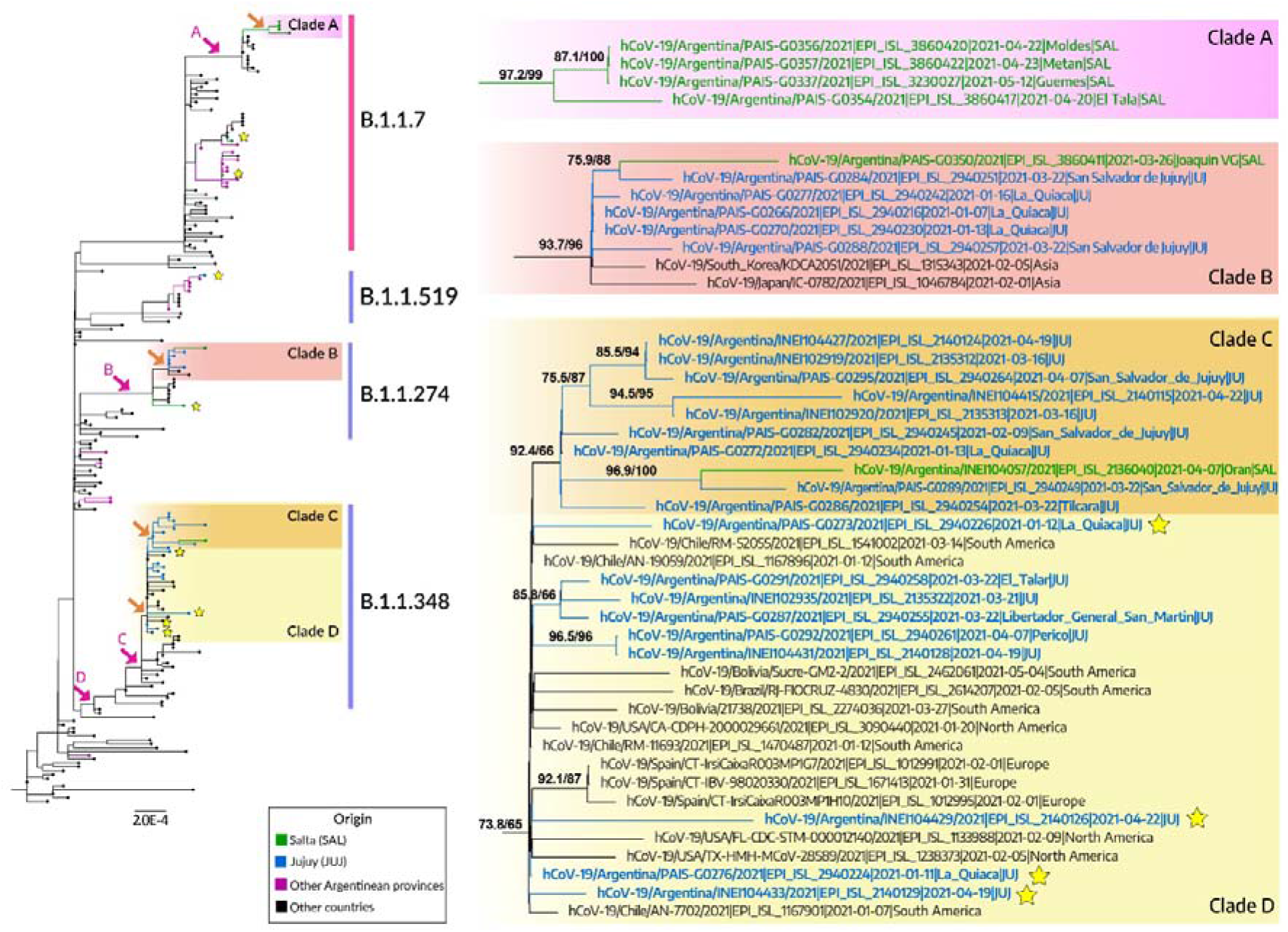
Phylogenetic tree of SARS-CoV-2 whole-genome sequences of lineage B.1.1 and its descendants. The tree was rooted between lineage A and B. Only groups with Northwestern Argentinean sequences are shown and indicated by orange arrows. The yellow stars indicate sporadic cases. The SH-like/UFB values for the relevant groups are indicated. Branches and tips are coloured by province. The scale indicates the number of substitutions per site. The pink arrows indicate the context groups used to analyse the descriptive mutations of some clusters that are detailed in **Table 2**.

#### VOI Lambda (lineage C.37)

18 sequences were assigned to the C.37 lineage: ten sequences from the province of Salta, six from Jujuy, and two from Tucumán. These sequences were collected during April and May 2021 and phylogenetic analysis determined eight introductions in the NWA region: five originating monophyletic groups and three associated with sporadic cases.

The SARS-CoV-2 genomes belonging to the Provinces of Salta and Jujuy showed monophyletic groups **(Figure 7, clades B and C)** that suggest local transmission and diversification of the Lambda variant in these provinces, but introductions in the Province of Tucumán, by contrast, corresponded to sporadic cases.

**Figure 7.**
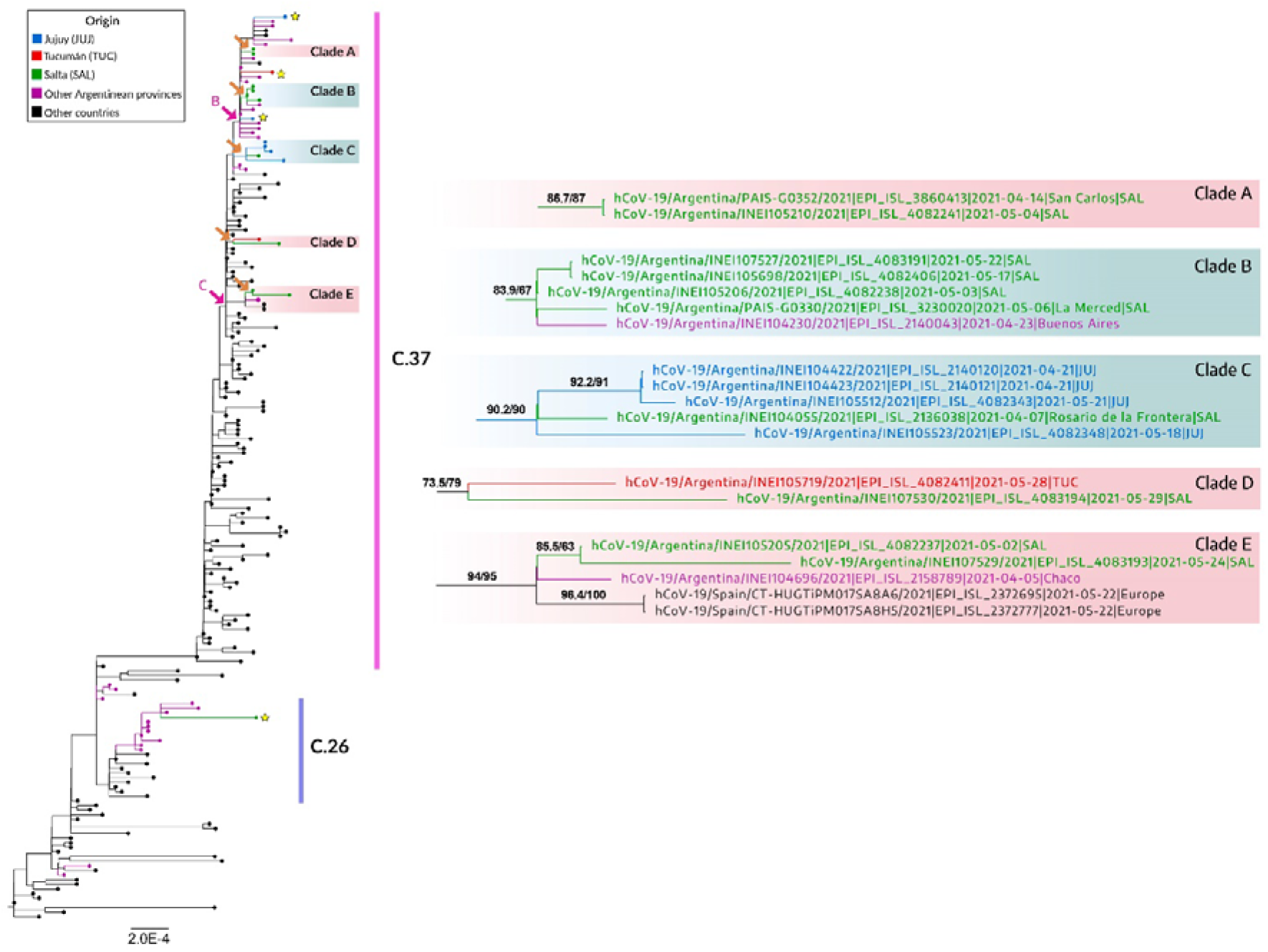
Phylogenetic tree of SARS-CoV-2 whole-genome sequences of Lambda (lineage C.37). Tree was rooted between lineages A and B. Only groups with Northwestern Argentinean sequences are shown and indicated by orange arrows. The yellow stars indicate sporadic cases. The SH-like/UFB values for the relevant groups are indicated. Branches and tips are coloured by province. The scale indicates the number of substitutions per site. The pink arrows indicate the context groups used to analyse the descriptive mutations of clusters B and C, which are detailed in **Table 2**.

#### VOI Zeta (lineage P.2)

Four genomes belonging to this lineage were detected during the early second wave (March-May 2021). Two of them belonged to the Province of Salta and formed a cluster along with sequences from Buenos Aires **(Figure 5, clade F)**. The other sequences corresponded to sporadic introductions without a phylogenetic relationship with other sequences from the NWA region.

#### VOI Epsilon (lineage B.1.427)

Nine sequences from the Province of Tucumán were assigned to this lineage, which was originally detected in California, the United States. The genomes were grouped into two clusters **(Figure 8, clades A and B)**, indicating that there were local transmission chains between March and May 2021.

**Figure 8.**
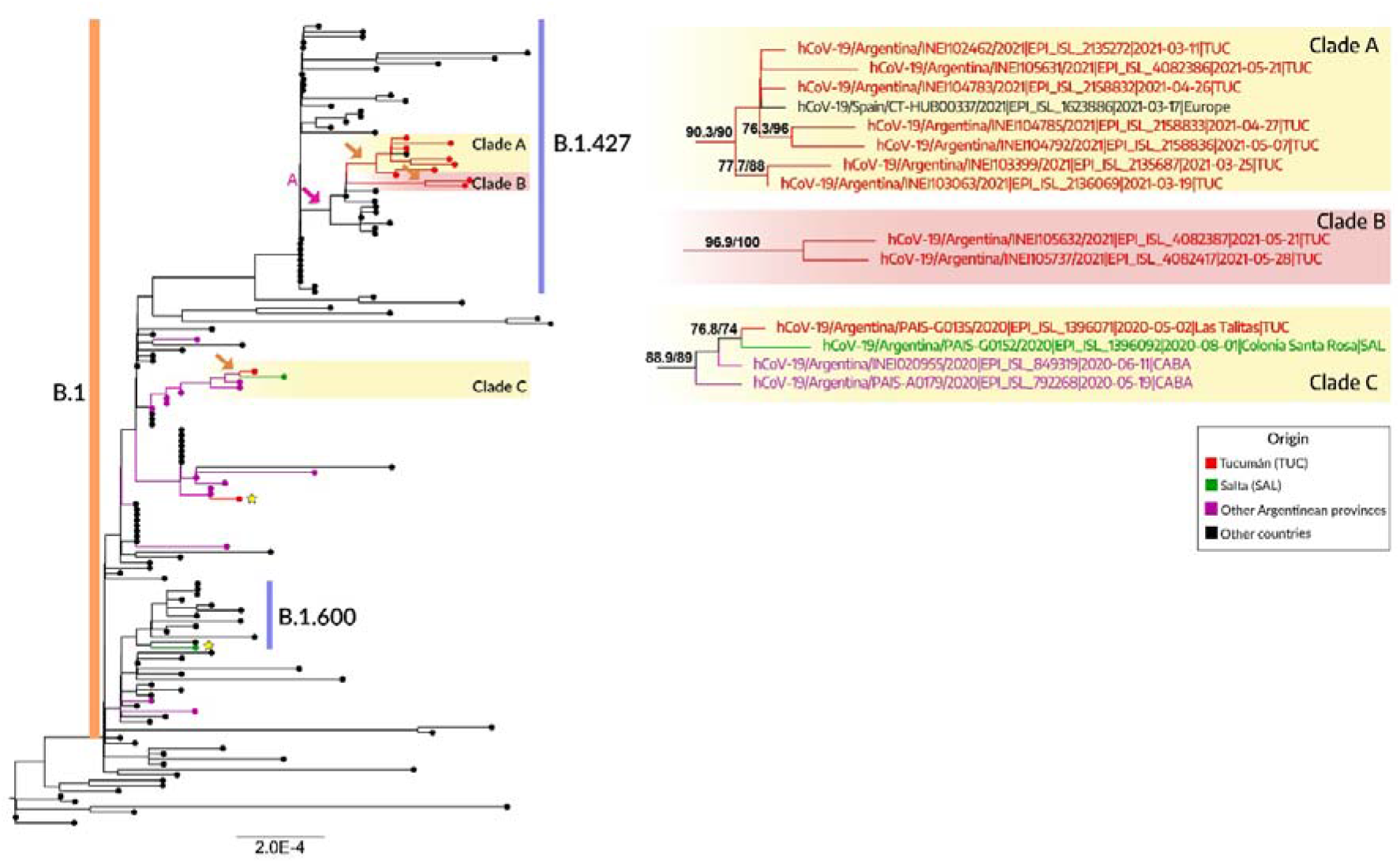
Phylogenetic tree of SARS-CoV-2 whole-genome sequences of Epsilon (lineage B.1.427). The tree was rooted between lineages A and B. Only groups with Northwestern Argentinean sequences are shown and indicated by orange arrows. The yellow stars indicate sporadic cases. The SH-like/UFB values for the relevant groups are indicated. Branches and tips are coloured by province. The scale indicates the number of substitutions per site. The pink arrow indicates the context group used to analyse the descriptive mutations of cluster A, which is detailed in **Table 2**.

#### Lineage A.2.5

Nine genomes from NWA (Salta and Tucumán) were assigned to this lineage and were grouped into a monophyletic group (Figure 9, Clade A) with sequences from other Argentinean provinces (Córdoba and Catamarca) and Chile. Internal diversification with three subclades can also be observed.

**Figure 9.**
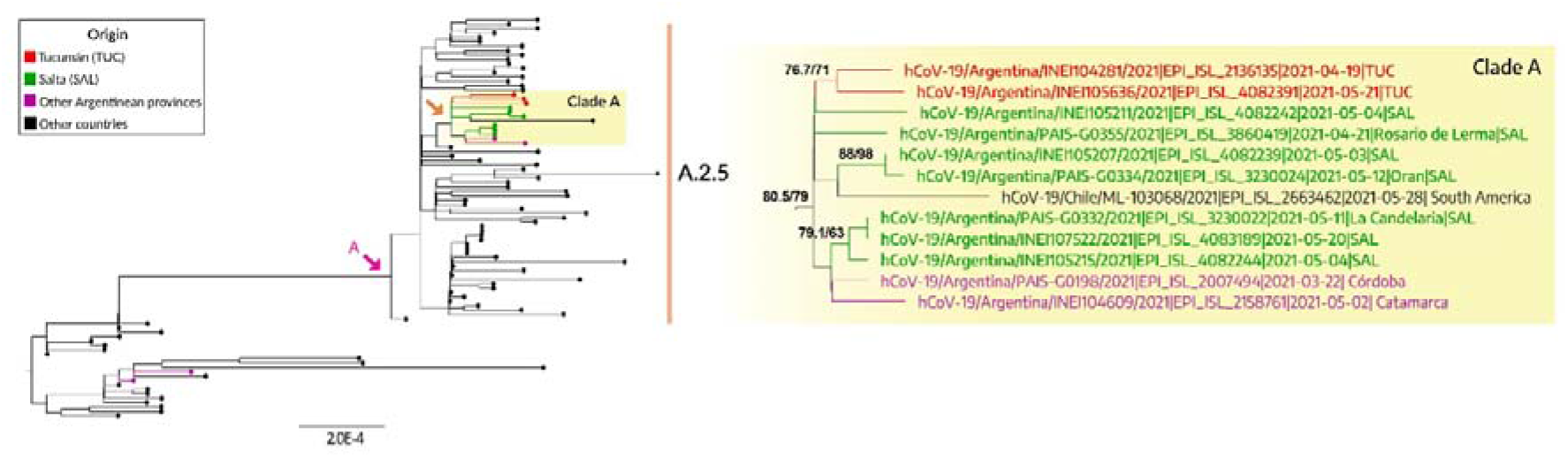
Phylogenetic tree of SARS-CoV-2 whole-genome sequences of lineage A.2.5. Tree was rooted between lineages A and B. Only groups with Northwestern Argentinean sequences are shown. The SH-like/UFB values for the relevant groups are indicated. Branches and tips are coloured by province. The scale indicates the number of substitutions per site. The pink arrow indicates the context group used to analyse the descriptive mutations of cluster A, which is detailed in **Table 2**.

### 3.4. Phylodynamics of Argentinean lineages in the NWA region

Phylodynamic analyses were performed on lineages B.1.499 and N.5, the two main Argentinean lineages that circulated in the first wave and the early second wave in the NWA region.

Lineage B.1.499 showed a substitution rate of 5.6 × 10^−4^ substitutions per site per year (s/s/y) (HPD95%= 4.5 × 10^−4^ - 6.9 × 10^−4^) and the date of its most recent common ancestor was estimated to be February 12, 2020 (HPD95%= January 8 – March 9). In addition, the demographic reconstruction of the lineage showed an increase in the effective number of infections in EW22/2020 (end of May) that persisted until a maximum in EW45/2020 (beginning of November) and decreased since then **(Figure 10)**.

**Figure 10.**
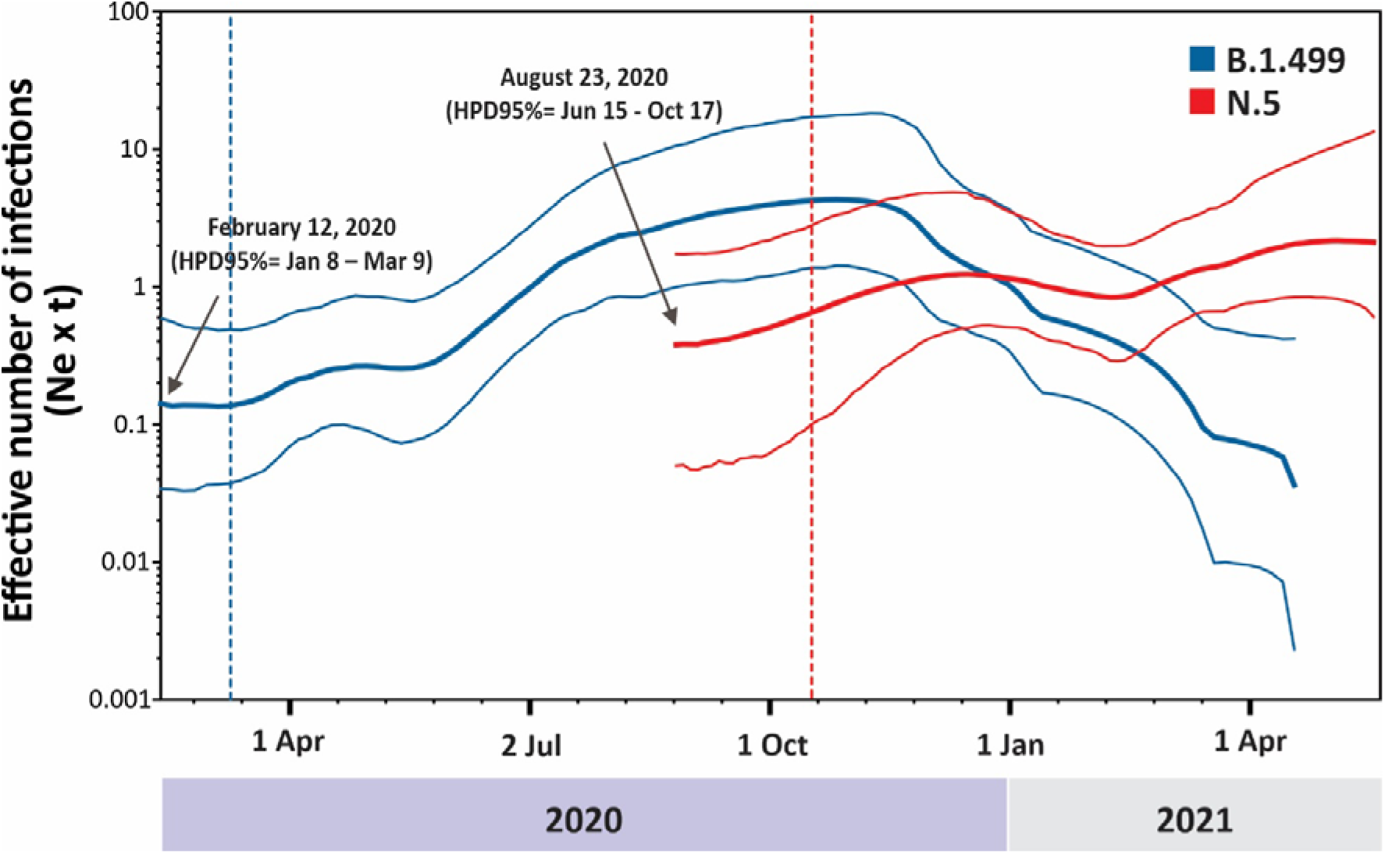
Demographic reconstruction of lineages B.1.499 and N.5. Estimations of the effective number of infections (inferred from the effective population size multiplied by the generation time, Ne × τ) over time from both lineages are superimposed. Mean values are shown as tick lines and HPD95% values (upper and lower), as thin lines. Dashed lines represent the upper HP95% interval for the time to the most recent common ancestor estimated for each lineage (indicated with arrows).

Lineage N.5 showed a substitution rate of 7.9 × 10^−4^ s/s/y (HPD95%= 5.6 × 10^−4^ - 1.0 × 10^−3^) and a most recent common ancestor dated on August 23, 2020 (HPD95%= June 15 - October 17), while its demographic reconstruction showed no substantial changes, only a trend of a slight increase over time **(Figure 10)**.

### 3.5. Mutations

Genomes sequenced obtained from NWA were compared with the Wuhan-Hu-4 reference sequence (*GISAID: EPI_ISL_402124*) to look for mutations in the Spike gene in the lineages with greater predominance in this study (Figure 11). In addition, characteristic mutations of VOCs were specially analyzed and were mostly found in single sequences but not in monophyletic groups as signature markers **(Table 1)**.

**Figure 11.**
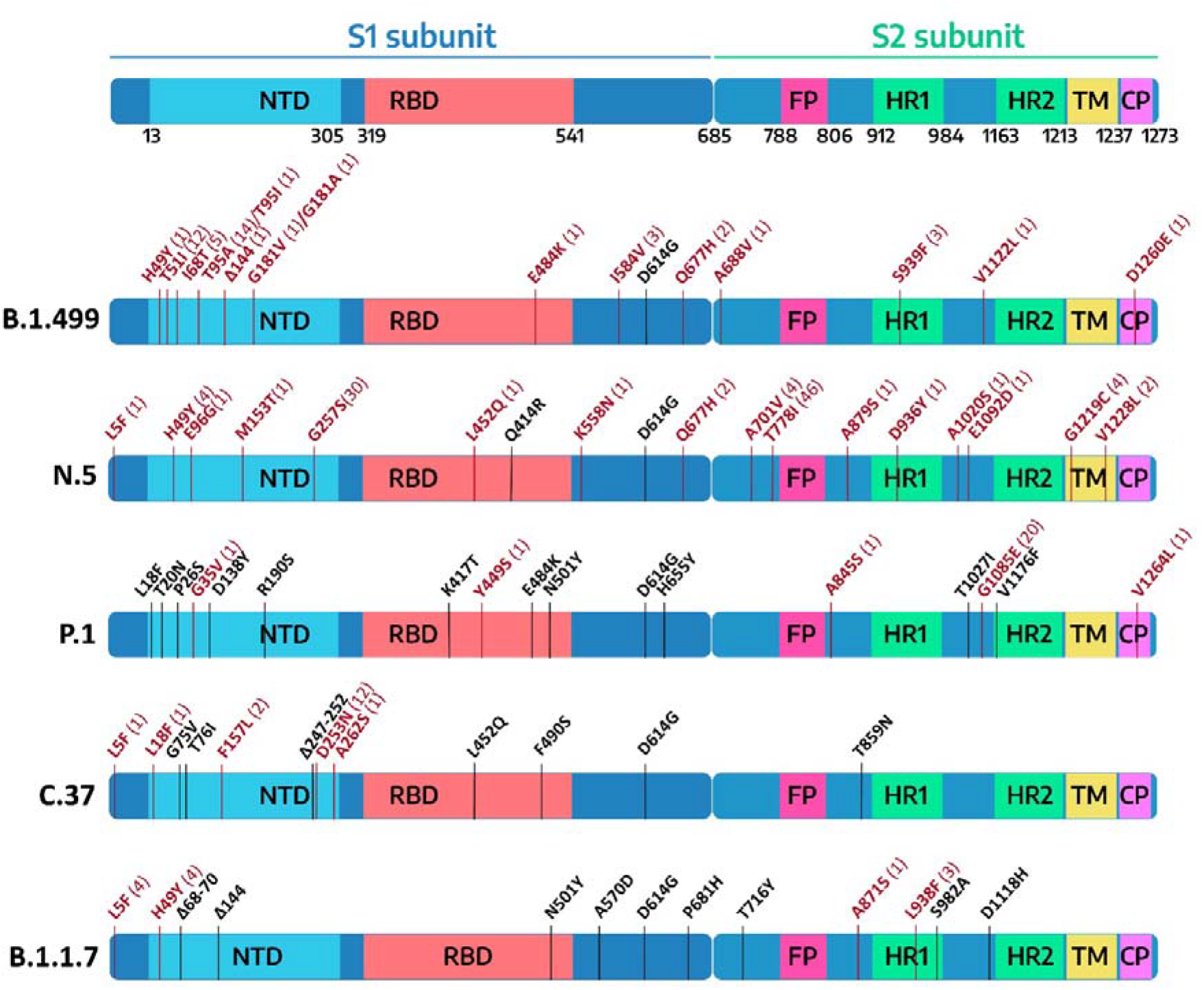
Schematic representation of amino acid changes in the Spike region found in most relevant lineages found in this study: B.1.499, N.5, P.1 (VOC Gamma), C.37 (VOI Lambda), B.1.1.7 (VOC Alpha), indicating key regions: NTD: N-terminal domain; RBD: Ribosomal binding domain; FP: Fusion peptide; HR1: Heptad repeat 1; HR2: Heptad repeat 2; TM: Transmembrane domain; CP: Cytoplasmic peptide. The mutations in black color correspond to the characteristic changes reported for the lineage and the mutations in red color to changes found for each lineage in this work. The number in parentheses indicates the number of sequences in which each mutation appeared.

**Table 1.** Significant mutations and their properties in the Spike protein of Northwestern Argentinean sequences from predominant lineages during the first and the early second wave.

| Mutation | GISAID ID | Lineage | VOC/VOI | Biological properties |
| --- | --- | --- | --- | --- |
| **E484K + Δ144** | EPI_ISL_2940196 | B.1.499 | **E484K:** Beta, Gamma, Iota, Theta, Zeta and Alpha.<br>**Δ144:** Alpha, Omicron and Eta. | **E484K:** Associated with an increment of the immunological resistance to neutralizing elicited by monoclonal and human serum antibodies (Zhou and Wang 2021).<br>**Δ144:** Resistance to several NTD-binding neutralizing mAbs but does not appear to reduce the neutralizing activity of plasma from convalescent or vaccinated patients (Li *et al.* 2020; McCallum *et al.* 2021; Wang *et al.* 2021). |
| **Q677H** | EPI_ISL_2940202,<br>EPI_ISL_2940198 | B.1.499 |  | Increases viral infectivity and syncytia formation and enhances resistance to neutralization by variants with RBD mutations (Zeng *et al.* 2021). |
|  | EPI_ISL_2136033,<br>EPI_ISL_4082236 | N.5 |  |  |
| **L452R** | EPI_ISL_2135314 | N.5 | Delta, Epsilon, Kappa and Omicron (lineages BA.4 and BA.5) | Associated with low-level reductions in susceptibility of convalescent and vaccine plasma samples (Ferreira *et al.* 2021; Greaney *et al.* 2021). |
| **Q675H** | EPI_ISL_4003459 | C.26 | P.6 | Improves proteolytic exposure of SARS-CoV-2 Spike in the Furin Binding Pocket (Bertelli *et al.* 2021). |
| **L18F** | EPI_ISL_4082302 | C.37 | Beta and Gamma | Associated with reduced susceptibility to several NTD-binding mAbs but by itself does not appear to reduce CP or VP susceptibility (McCallum *et al.* 2021; Wang *et al.* 2021). |

In addition, we analyzed signature mutations in **19 phylogenetic groups** containing Argentinean sequences in different lineages. In total, 42 group-defining mutations were observed: 22 were non-synonymous mutations, 19 were synonymous mutations, and one of them corresponded to a substitution in the non-coding 5’ region of the viral genome **(Table 2)**.

**Table 2.** A detailed description of mutations harboured by the monophyletic groups in lineages of the first and the early second wave in the NWA.

| Lineage | Clade name<sup>a</sup> | Nucleotide mutation | Gene/region | Effect | Amino Acid change<sup>b</sup> |
| --- | --- | --- | --- | --- | --- |
| A.2.5 | A | G443A | ORF1a/nsp1 | Non-synonymous | V60I |
|  |  | C27143T | M | Synonymous | N207- |
| B.1.427 | A | C14262T | ORF1b/nsp12 | Synonymous | D274- |
|  |  | T15741C | ORF1b/nsp12 | Synonymous | N767- |
| B.1.1.7 | A | T2169G | ORF1a/nsp2 | Non-synonymous | F455C |
|  |  | G14316A | ORF1b/nsp12 | Synonymous | Q292- |
|  |  | G16221A | ORF1b/nsp12 | Synonymous | P927- |
|  |  | G19684T | ORF1b/nsp15 | Non-synonymous | V22L |
|  |  | C21707T | Spike | Non-synonymous | H49Y<sup>c</sup> |
| B.1.1.274 | B | C23604A | Spike | Non-synonymous | P681H<sup>d</sup> |
|  |  | A25998T | ORF3a | Synonymous | V202- |
|  |  | C29375T | N | Non-synonymous | P368S |
| B.1.1.348 | C | G5572T | ORF1a/nsp3 | Non-synonymous | M951I<br>M951V |
|  | D | T22679C | Spike | Non-synonymous | S373P |
| B.1.499 | A | C7279T | ORF1a/nsp3 | Synonymous | F1520- |
|  |  | **C7528T** | ORF1a/nsp3 | Synonymous | **V1603-** |
|  | **B** | **A23989T** | Spike | Synonymous | **P809-** |
|  |  | **T27534C** | ORF7a | Synonymous | **H47-** |
|  | **D** | **C11750T** | ORF1a/nsp6 | Non-synonymous | **L260F** |
|  | **E** | **A8031G** | ORF1a/nsp3 | Non-synonymous | **K1771R** |
|  |  | **C21714T** | Spike | Non-synonymous | **T51I** |
|  | **F** | **C2334T** | ORF1a/nsp2 | Non-synonymous | **A510V** |
|  |  | **C28054A** | ORF8 | Non-synonymous | **S54X** |
|  | **G** | **A21845G** | Spike | Non-synonymous | **T95A** |
| **C.37** | **B** | **A27486G** | ORF7a | Synonymous | **L31-** |
|  | **C** | **T63C** | non coding | ----- | ----- |
|  |  | **C3130T** | ORF1a/nsp3 | Synonymous | **Y137-** |
| **N.5** | **A** | **C7765T** | ORF1a/nsp3 | Synonymous | **S1682-** |
|  | **B** | **C12049T** | ORF1a/nsp7 | Synonymous | **N69-** |
|  | **D** | **G22801T** | Spike | Synonymous | **G413-** |
|  | **I** | **A454G** | ORF1a/nsp1 | Synonymous | **Q63-** |
|  |  | **C2358T** | ORF1a/nsp2 | Non-synonymous | **A518V** |
|  |  | **C4965T** | ORF1a/nsp3 | Non-synonymous | **T749I** |
|  |  | **G5470A** | ORF1a/nsp3 | Synonymous | **L91-** |
|  |  | **A5828G** | ORF1a/nsp3 | Non-synonymous | **K1037E** |
|  |  | **G6446A** | ORF1a/nsp3 | Non-synonymous | **V1243I** |
|  |  | **T14727C** | ORF1b/nsp12 | Synonymous | **F429-** |
|  |  | **C16332T** | ORF1b/nsp13 | Synonymous | **D32-** |
|  |  | **C23664T** | Spike | Non-synonymous | **A701V<sup>d</sup>** |
| **P.1** | **A** | **G24816A** | Spike | Non-synonymous | **G1085E** |

<sup>a</sup> The names of the clades correspond to those mentioned in the figures of each lineage.
<sup>b</sup> Dash (-) represents amino acid maintenance (synonymous change).
<sup>c</sup> H49Y is related to greater stabilizing interaction of the ACE2-Spike complex (Aljindan *et al.* 2021).
<sup>d</sup> P681H and A701V do not play a role in viral infectivity and neutralization potential. However, when in conjunction with the RBD-N501Y mutation, viral infectivity is enhanced (Kuzmina *et al.* 2022).

Non-synonymous mutations were found particularly in the ORF1a (n= 12) and S (n= 7) genes and, to a lesser extent, in the ORF1b, N and ORF8 genes with one mutation recorded in each of them. In the case of the synonymous mutations, they were mainly detected in the ORF1a (n= 6) and ORF1b (n= 6) genes and were also observed in the S (n= 2), ORF7a (n= 2), M (n= 1) and ORF3a (n= 1) genes.

In the **Spike, the non-synonymous mutations** were: H49Y (C21707), P681H (C23604A), S373P (T22679C), T51I (C21714T), T95A (A21845G), A701V (C23664T), G1085E (G24816A). Some of these mutations are of particular interest.

The H49Y mutation is one of the changes that characterize the clade A of VOC Alpha from the NWA **(Figure 6, clade A)** that also appeared in the lineage B.1.499 (GISAID: EPI_ISL_1396094). The mutation P681H, characteristic of the VOC Alpha, VOC Omicron and VOI Mu, was found in clade B (lineage B.1.1.274) **(Figure 6, clade B)**. Finally, the **A701V** mutation, observed in the VOC Beta and the VOI Iota, was also associated with clade I of lineage N.5 (a clade with nine defining-group mutations) **(Figure 4, clade I)**.

## 4. Discussion

In this work, we analyzed the evolution of SARS-CoV-2 during the first year of the COVID-19 pandemic in Northwestern Argentina, covering the first and the early second wave in the region.

The first wave was dominated by the lineage B.1.499, an almost exclusive Argentinean lineage. Its common ancestor was estimated to be dated in mid-February 2020, although the first sample in the country was detected at the beginning of April 2020, in a sample from NWA. The demographic reconstruction of lineage B.1.499 showed an increase in the effective number of infections that matched the reported cases in the National Surveillance System during the first wave in the NWA, evidencing that this was the lineage that mainly impulses this wave in the region.

The second most prevalent lineage in the first wave in the NWA was the lineage N.5, derived from B.1.1.33, a Brazilian lineage that dispersed mainly into South American countries in the first months of the pandemics (Resende *et al*. 2021). In this case, the phylodynamic analysis showed a common ancestor diversifying in late-August 2020, much earlier than the first detection in the NWA region (beginning of January 2021), suggesting that the viruses that reached the NWA originated from different transmission chains previously established in other regions of the country.

On the other hand, it was suggested that the emergence of the VOCs Alpha to Delta was driven by an acceleration of the substitution rate in their branch stems, with estimations rounding an acceleration of 4-fold, with a mean “background” (non-VOC) substitution rate of 5.8 × 10^−4^ s/s/y and a rate of 2.5 × 10^−3^ s/s/y for the VOC stems (Tay *et al*. 2022). These results encourage the monitoring of the molecular evolution of SARS-CoV-2 as a tool to understand the context in which a lineage of epidemiological interest can emerge. In this framework, the analysis of the evolutionary dynamics of the two most prevalent lineages in NWA also revealed that the substitution rate of lineage N.5 (7.9 × 10^−4^ s/s/y) was a ∼40% faster than that of lineage B.1.499 (5.9 × 10^−4^ s/s/y), although both are in the same order of magnitude than other non-VOC lineages (Tay *et al*. 2022). These results showed the importance of analyzing substitution rates separately for different lineages, whenever possible, rather than assuming a fixed rate for all, to improve the understanding of the evolutionary dynamics of SARS-CoV-2 lineages.

It should be noted that for the phylodynamic analyses included in this work, a selection of sequences belonging to the lineages B.1.499 and N.5 from the NWA was made to focus on the evolutionary dynamics of viruses from this region. However, to deepen the study of the behavior of the lineages, sequences from the entire country and a longer period would be necessary.

Some few mutations associated with a biological characteristic of importance were observed as signatures markers of the phylogenetic groups established in Northwestern Argentina, and even though positive selection cannot be ruled out, their defining-groups mutations more probably reflect neutral evolution, a major characteristic of the first wave all around the world (Dearlove *et al*. 2020). Like other first-wave lineages, except for the D614G in Spike (Korber *et al*. 2020), the P323L in the viral RNA-dependent RNA polymerase protein (Garvin *et al*. 2020) and other sporadic mutations in Spike, they did not present mutations associated with adaptation to the host as lineages markers (as VOC did) (Martin *et al*. 2021; Peacock *et al*. 2021). However, single sequences in non-VOC lineages did present mutations associated with VOCs, showing that many of these mutations could emerge from circulation in the general population.

During the early second wave, the first-wave lineages were displaced by the introduction of VOCs (Alpha, Gamma), VOIs (Lambda, Zeta, Epsilon) and other lineages with more limited distribution.

It is worth noting that VOC Alpha did not become established in the NWA, as it was observed in most other regions outside of South America (Hodcroft 2021). On the contrary, VOC Gamma (lineage P.1 and derivatives) and VOI Lambda (lineage C.37) were of importance in the second wave of the COVID-19 pandemic in South America, particularly in Argentina (Faria *et al*. 2021; Romero *et al*. 2021; Torres *et al*. 2021). Several introductions were observed for these variants in the NWA, with sequences associated with samples from other regions of the country or neighboring countries (Brazil and Chile). However, the limited information on molecular surveillance of SARS-CoV-2 in Bolivia and Paraguay does not allow for an evaluation of the complete landscape of the region in the studied period.

Within the NWA region, and despite being composed of three provinces that share borders, it was shown that different routes of viral dispersion impact the epidemiology of the region. For instance, Salta and Jujuy showed evidence of common transmission chains in different lineages, whereas sequences from Tucumán associated with sequences from other provinces in the country, suggesting that internal movement due to commercial and touristic routes impact differentially on the provinces.

Finally, this study contributed to the knowledge about the evolution of SARS-CoV-2 in a pre-vaccination and without post-exposure immunization period, gaining relevance for the understanding of the evolutionary behavior of SARS-CoV-2 in a naive environment. These analyses can establish a background level for new analyses on these or other SARS-CoV-2 lineages in later periods when other factors that may have an impact on the viral evolution and emergence of variants appeared, such as post-exposure immunity, vaccination, waning immunity, among others.

## 5. Conflict of Interest

The authors declare that the research was conducted in the absence of any commercial or financial relationships that could be construed as a potential conflict of interest.

## 6. Author Contributions

RZ performed analyses, analyzed data, and co-wrote the article. AC, FF, NM, HD performed experiments, analyzed data, and provided critical revision of the article. MS, AZ, GR, DC, GA, MC, AF, CM, FV, CD, VR, EL, VL, PC, FA, CC collected samples, performed experiments, and provided data. CT designed analyses, analyzed data, and co-wrote the article. MV analyzed data, provided critical revision of the article, and coordinated the project.

## 7. Acknowledgement

This work was supported by Proyecto IP COVID-19 N°08 (Ministerio de Ciencia, Tecnología e Innovación, Argentina) and Focem COF 03/11 Covid-19 (Fondo para la Convergencia Estructural del MERCOSUR). Funding had no role in the study design, collection, analysis, or interpretation of data, in the writing or in the decision to submit the article for publication.

## Notes

### Competing Interest Statement

The authors have declared no competing interest.

