## Supplementary material for "Evolution of SARS-CoV-2 during the first year of the COVID-19 pandemic in Northwestern Argentina": Table S2

### SUPPLEMENTAL TABLE

**Data Availability**

| | |
|---|---|
| GISAID Identifier: | EPI_SET_20220615tq |
| DOI: | https://doi.org/10.55876/gis8.220615tq |

All genome sequences and associated metadata in this dataset are published in GISAID's EpiCoV database. To view the contributors of each individual sequence with details such as accession number, Virus name, Collection date, Originating Lab and Submitting Lab and the list of Authors, visit https://doi.org/10.55876/gis8.220615tq

**Data Snapshot**

- EPI_SET_20220615tq is composed of 1,302 individual genome sequences.
- The collection dates range from 2020-01-05 to 2021-07-20;
- Data were collected in 46 countries and territories;
- All sequences in this dataset are compared relative to hCoV-19/Wuhan/WIV04/2019 (WIV04), the official reference sequence employed by GISAID (EPI_ISL_402124). Learn more at https://gisaid.org/WIV04.
